# Genetic potential for aerobic respiration and denitrification in globally distributed respiratory endosymbionts

**DOI:** 10.1101/2024.02.19.580942

**Authors:** Daan R. Speth, Linus M. Zeller, Jon S. Graf, Will A. Overholt, Kirsten Küsel, Jana Milucka

## Abstract

The endosymbiont *Candidatus* Azoamicus ciliaticola generates ATP for its eukaryotic host, an anaerobic ciliate of the Plagiopylea class, fulfilling a function analogous to mitochondria in other eukaryotic cells. The discovery of this respiratory endosymbiosis has major implications for both evolutionary history and ecology of microbial eukaryotes. However, with only a single species described, knowledge of its environmental distribution and diversity is limited. Here we report four new complete, circular metagenome assembled genomes (cMAGs) representing respiratory endosymbionts inhabiting groundwater in California, Ohio, and Germany. These cMAGs form two lineages comprising a monophyletic clade within the uncharacterized gammaproteobacterial order UBA6186, enabling evolutionary analysis of their key protein complexes. Strikingly, all four novel cMAGs encode a cytochrome *cbb_3_* oxidase, which indicates the capacity for both aerobic and anaerobic respiration. Accordingly, we detect these respiratory endosymbionts in diverse habitats worldwide, thus significantly expanding the ecological scope of this respiratory symbiosis.

## Main

Host beneficial endosymbionts (HBEs) are widespread in eukaryotes, and often evolve from a parasitic or pathogenic ancestor (McCutcheon, Boyd, and Dale 2019). The best studied HBEs are nutritional symbionts that provide their hosts with essential molecules (such as vitamins or cofactors), which the hosts cannot synthesize or obtain from their diet (Douglas 2003; Moran et al. 2003). Another example are defensive endosymbionts that synthesize compounds that protect the host or its offspring against predators or pathogens (Brownlie and Johnson 2009; Van Arnam, Currie, and Clardy 2018). The recently discovered protist endosymbiont *Ca.* A. ciliaticola fulfills yet another function by generating ATP using a denitrifying respiratory chain and supplying it to its ciliate host (Graf et al. 2021). The Plagiopylean ciliate host of *Ca.* A. ciliaticola appears to have lost its mitochondria, and is thus incapable of (aerobic) respiration, although it might still be able to generate ATP through fermentation using hydrogenosomes, or substrate level phosphorylation in the cytoplasm (Embley et al. 1997; Graf et al. 2021). *Ca.* A. ciliaticola lacks the capability for aerobic respiration and is obligately anaerobic, restricting the ecological niche of its ciliate host to permanently anoxic habitats, such as the hypolimnion of a meromictic lake. Interestingly, virtually all organisms capable of denitrification are facultative anaerobes (Chen and Strous 2013; Kuypers, Marchant, and Kartal 2018), and therefore it was hypothesized that *Ca*. A. ciliaticola evolved from a predecessor that was capable of both aerobic respiration and denitrification (Graf et al. 2021). To investigate this possibility, we searched for genomes of endosymbionts related to *Ca.* A. ciliaticola in publicly available environmental metagenomic sequencing datasets.

## Results

### Discovery and genomic features of novel endosymbiont genomes

We used the *tlcA* gene, which encodes for an ATP/ADP transporter that is crucial for the ATP supplying function of *Ca.* A. ciliaticola, to identify putative respiratory endosymbionts. Using this strategy, we recovered one new complete cMAG originating from groundwater samples taken in California (He et al. 2021) and assembled three new complete cMAGs from groundwater samples of a carbonate-rock-fracture aquifer taken in Germany (Overholt et al. 2022; Küsel et al. 2016) and Ohio (2 cMAGs) (Danczak et al. 2019). All four new cMAGs share genomic features characteristic of obligate endosymbionts, including extreme genome reduction (284-373 kb genome size), low GC content (20-26 %) and a high protein coding density (91-92 %) (Figure 1, Extended data table 1). Additionally, all four cMAGs contain a complete respiratory denitrification pathway (Figure 1, Supplemental table S1), but have undergone extensive loss of genes for the biosynthesis of vitamins, amino acids, and other essential metabolites (Supplemental table S2). These genomic similarities strongly suggest that, like *Ca*. A ciliaticola, these cMAGs represent obligate respiratory endosymbionts that perform denitrification and provide ATP for their hosts.

**Figure 1.**
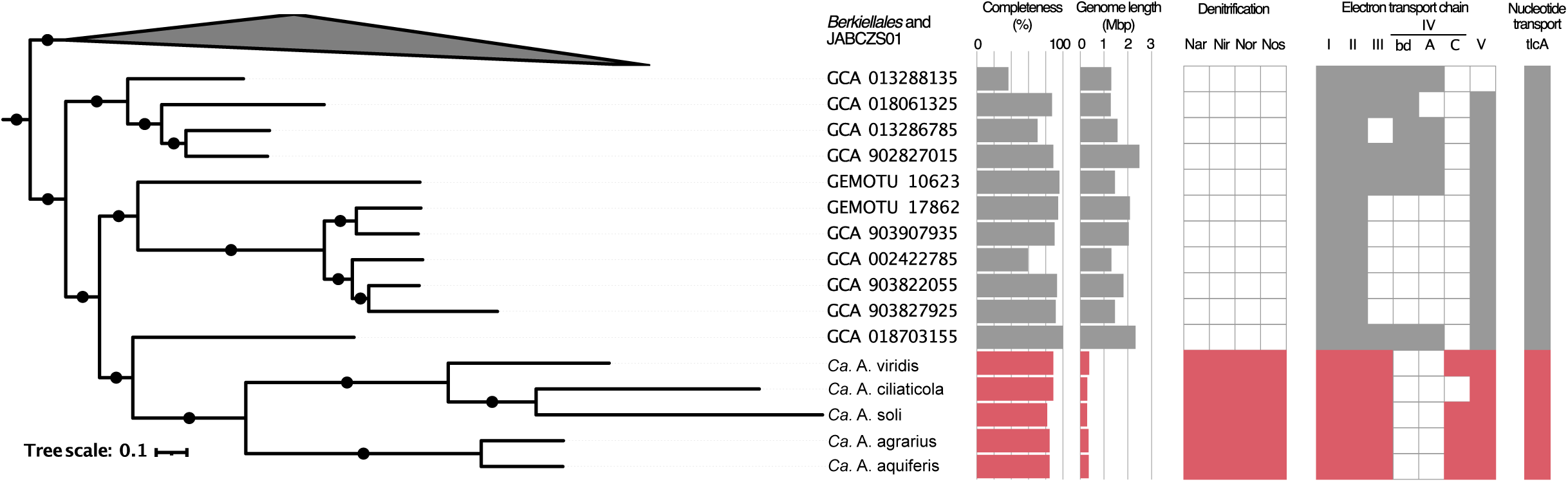
Phylogeny and selected genomic features of the UBA6186 order. Concatenated marker gene phylogeny of the gammaproteobacterial orders Berkiellales, JABCZS01 and UBA6186. The expanded section shows the UBA6186 order, including the Azoamicaceae. The tree annotations indicate, from left to right, genome size, MAG completeness as estimated by Anvi’o, presence/absence of the denitrification pathway, presence/absence of respiratory complexes NADH dehydrogenase (I), succinate dehydrogenase (II), bc_1_ complex (III), cytochrome bd oxidase (IV - bd), cytochrome caa_3_ oxidase (IV - A), cytochrome cbb_3_ oxidase (IV - C), and ATP synthase (V), and presence of at least one copy of the tlcA gene encoding an ATP/ADP transporter. The genomic features of Azoamicaceae are indicated in red, while the features of the other UBA6186 genomes are shown in grey. Sequences were obtained from genomes included in the genome taxonomy database and global catalog of earth’s microbiomes databases. Black circles on the branches indicate bootstrap values higher 90%.

The four new cMAGs (hereafter: genomes) form a monophyletic clade with *Ca.* A. ciliaticola, within the UBA6186 order of the Gammaproteobacteria (Figure 1, Extended Data Figure 1, Supplemental Text). The groundwater genomes belong to two distinct clades, with the two ‘Ohio’ genomes forming a lineage with the lacustrine *Ca.* A. ciliaticola, and the ‘California’ and ‘Germany’ genomes forming a novel lineage (Figure 1). Based on the pairwise average amino acid identity (AAI) and average nucleotide identity (ANI) values (Jain et al. 2018; Barco et al. 2020) between the five genomes (Supplemental table S3) we propose that they represent five species grouped in two genera within the *Candidatus* Azoamicaceae family (hereafter: Azoamicaceae). The two genera are the previously established *Candidatus* Azoamicus (hereafter: Azoamicus) and a novel genus for which we propose the name *Candidatus* Azosocius (hereafter: Azosocius; see Supplemental Text for discussion of boundaries for species and genera). We propose the names *Candidatus* Azoamicus viridis and *Candidatus* Azoamicus soli for the ‘Ohio’ genomes and *Candidatus* Azosocius agrarius and *Candidatus* Azosocius aquiferis for the ‘California’ and ‘Germany’ genomes, respectively. *Candidatus* Azosocius agrarius is the type species of the Azosocius genus.

The genomes of the Azosocius species are strikingly similar in size (352 and 353 kpb for *Ca.* A. agrarius and *Ca.* A. aquiferis, respectively) and they are larger than those of *Ca.* A. soli (284 kpb) and *Ca.* A. ciliaticola (293 kbp); however, the largest new genome belonged to *Ca.* A. viridis (374 kbp). A comparative genomic analysis of the five Azoamicaceae genomes revealed that their collective protein coding gene content amounts to only 470 genes (Figure 2a), of which 254 are shared between all five genomes (68 - 87 % of the protein coding gene complement of individual genomes). This conserved set includes the genes for an electron transport chain and ATP synthase (Complexes I, II, III, V), a complete denitrification pathway for reduction of nitrate to dinitrogen gas, ATP/ADP transporter, iron sulfur cluster biosynthesis, molybdopterin cofactor biosynthesis, and cellular information processing (Figure 2d, Supplemental table S1). 43 additional genes are present in multiple (2-4) genomes across both Azoamicus and Azosocius lineages (Figure 2a), indicating ongoing loss of genes after the lineages diverged. In addition to the genes shared between the two lineages, there are 107 genes unique to Azoamicus and 66 genes unique to Azosocius (Figure 2b), as further discussed in the Supplementary Text.

**Figure 2.**
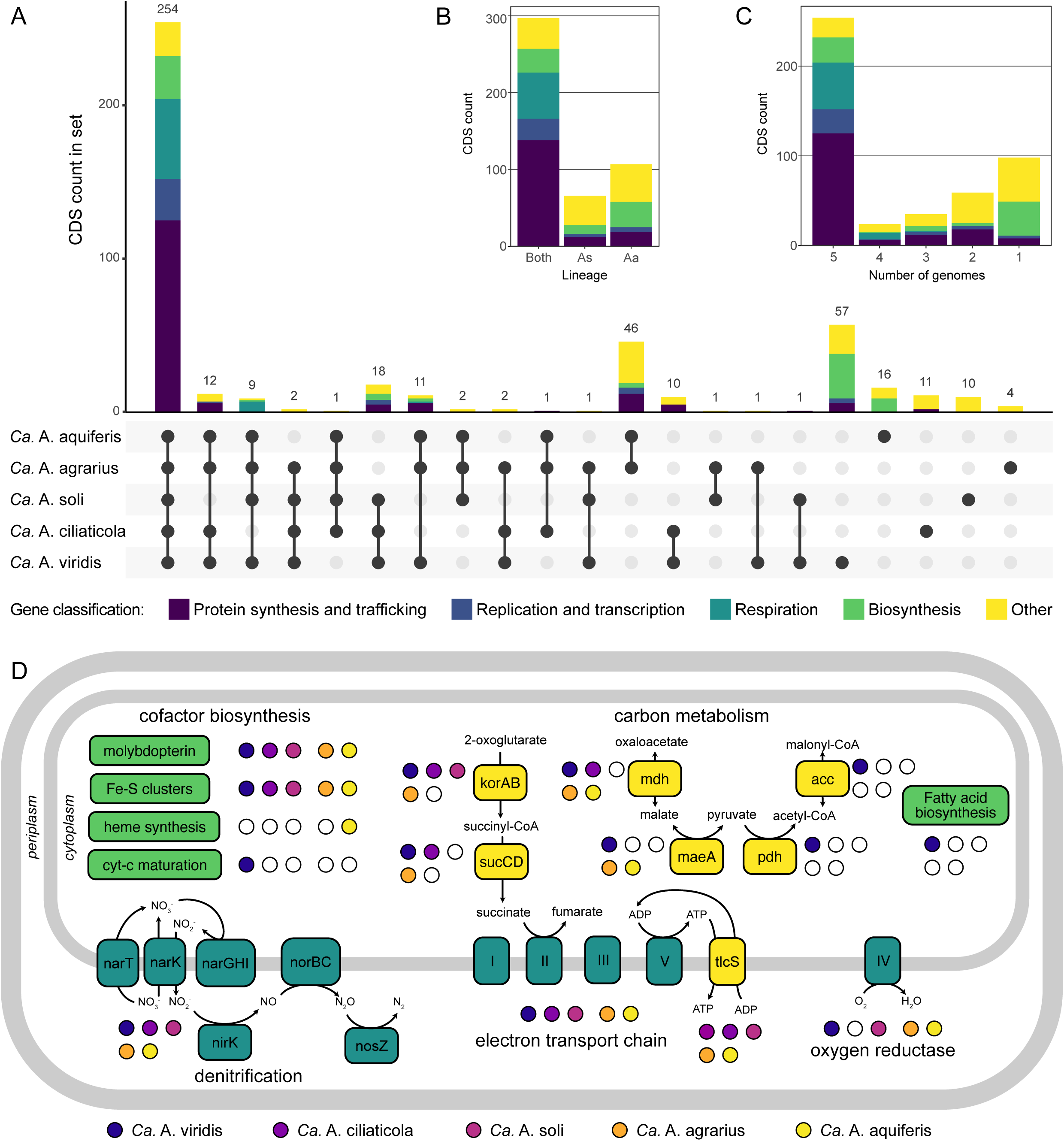
Gene distribution and pathway overview in the Azoamicaceae genomes. (A) UpSet plot showing the coding sequence (CDS) content shared between, or unique to, the individual Azoamicaceae genomes. Each vertical bar represents the number of genes in an intersecting set between the genomes. The filled circles connected by lines indicate which genomes contribute genes to the intersecting set. Sets are ordered by the number of genomes contributing, and then the number of genes in the set. Vertical bars are colored according to the broad classification of genes that can be found in supplemental table S1. Total coding sequence numbers for the genomes are Ca. A aquiferis 342; Ca. A agrarius 344; Ca. A soli 293; Ca. A ciliaticola 310; Ca. A viridis 372 (B) (inset) Distribution of the gene content shown in panel A, summarized by genus. Aa: Azoamicus, As: Azosocius. (C) (inset) Distribution of the CDS content shown in panel A summarized by the number of genomes containing each gene. (D) Key pathways in the Ca. Azoamiceae genomes that enable generation of ATP for the host cell by both denitrification and aerobic respiration. Boxes indicate protein complexes or pathways, with background shading corresponding to the categories in panels A - C. Filled and empty circles indicate the presence/absence of a complex or pathway in the genome. Roman numerals in the boxes representing the electron transport chain complexes indicate: (I) NADH dehydrogenase, (II) succinate dehydrogenase, (III) bc_1_ complex, (IV) cytochrome cbb_3_ oxidase, (V) ATP synthase.

### Presence and transcription of high affinity cytochrome c oxidase in groundwater Azoamicaceae

Notably, among the genes conserved in the four new genomes is an operon consisting of ccoNO(Q)P, encoding for a high affinity cytochrome *cbb*_3_ oxygen reductase (complex IV) (Pitcher and Watmough 2004). The *cbb*_3_ operon is complete in three of the four new genomes, only *Ca.* A. soli appears to have lost the *ccoQ* gene that encodes for the small non-catalytic subunit (Kohlstaedt et al. 2017). The cytochrome *cbb*_3_ oxygen reductases were initially thought to be restricted to Proteobacteria, but have since also been found in genomes of organisms from other bacterial phyla (Ducluzeau, Ouchane, and Nitschke 2008). In organisms encoding a cytochrome *cbb_3_*oxidase, small c_4_ and c_5_ cytochromes can form a branch point in the respiratory chain, allowing electrons from complex III to flow to either the *cbb_3_*oxidase during aerobic respiration, or a nitrite reductase during denitrification. All Azoamicaceae encode the diheme cytochrome c_4_ that acts as the natural electron donor to the cytochome *cbb*_3_ oxidase in *Vibrio cholerae* (Chang et al. 2010) and *Neisseria meningitidis* (Deeudom, Koomey, and Moir 2008), as well as the c_5_ monoheme cytochrome that shuttles electrons from complex III to nitrite reductase in *Neisseria meningitidis* (Deeudom, Koomey, and Moir 2008). These small cytochromes are also present in *Ca.* A. ciliaticola which lacks the *cbb_3_* oxidase, strongly suggesting that *Ca.* A. ciliaticola has lost its terminal oxidase secondarily. This is further supported by the presence of a *cbb_3_* oxidase in closely related *Ca.* A. viridis and *Ca.* A. soli genomes from the Azoamicus genus (Figure 1).

We confirmed the transcription of the *ccoNOQP* operon in *Ca.* A aquiferis using data from 18 groundwater metatranscriptome datasets of the Hainich Critical Zone Exploratory (CZE) (Wegner et al. 2019), from which the *Ca.* A aquiferis genome was assembled (Supplemental table S5). Sufficient reads matching *Ca.* A. aquiferis genes for analysis of genome wide transcription patterns (327 - 1138 reads) could be recovered from seven of these datasets, which represent two wells sampled at two time points. The overall observed transcription pattern was very similar in all seven datasets (Figure 3, Supplemental table S5). As previously observed for *Ca*. A. ciliaticola, the *nirK*, *norB*, and *nosZ* genes were amongst the highest transcribed genes for *Ca.* A aquiferis in all groundwater samples, with transcription of the other genes of the denitrification pathway lower but still detectable (Graf et al. 2021). The catalytic subunit of the cytochrome *cbb_3_*oxidase (*ccoN*) was also consistently highly transcribed, at a similar level as e.g. the *narG* gene (encoding for the catalytic subunit of nitrate reductase). The *ccoO* and *ccoP* subunits of the cytochrome *cbb_3_* oxidase were also highly transcribed in some but not all datasets (Figure 3, Supplemental Table S5). Interestingly, high *ccoNOP* gene transcription was detected in samples with markedly different dissolved oxygen concentrations (Küsel et al. 2016). Samples from well H5-2 originate from typically anoxic groundwater in low permeable marlstones and dissolved oxygen was not detected at the time of RNA sampling (Kohlhepp et al. 2017). In contrast, groundwater samples from the permeable fracture aquifer accessed at well H4-1 contained approximately 6 mg/L dissolved oxygen at the time of RNA sampling (Kohlhepp et al. 2017), which is typical for this sampling site (Küsel et al. 2016). The comparably high levels of transcription of the cytochrome *cbb*_3_ oxidase operon in oxic and anoxic groundwater at both timepoints (Figure 3) may indicate that the terminal oxidase is constitutively expressed in *Ca.* A. aquiferis, as previously observed for *cco1* of *Pseudomonas aeruginosa* (Kawakami et al. 2010).

**Figure 3.**
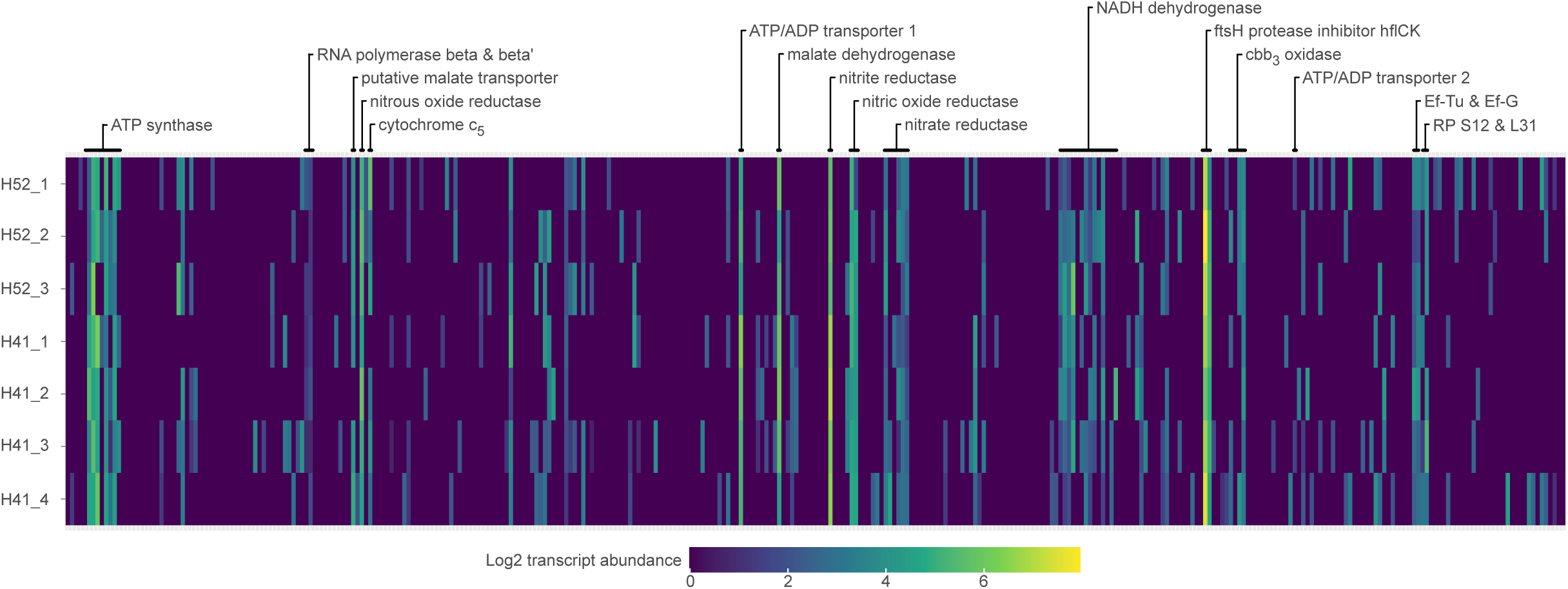
In situ gene expression of Ca. A. aquiferis genes in groundwater metatranscriptomes from Hainich CZE. Heatmap depicting the gene expression of 352 protein coding genes (columns) in the genome of Ca. A. aquiferis across seven metatranscriptome samples (rows) taken from anoxic groundwater of well H5-2 and oxic groundwater of well H4-1 at the Hainich critical zone exploratory (CZE) during two sampling campaigns. The genes are arranged in order according to their position in the Ca. A. aquiferis genome, with highly expressed genes and other selected genes annotated above the heatmap. The gene expression was normalized to “transcripts per thousand”, and then log_2_ transformed to highlight the genome-wide expression pattern. Ef-Tu: Elongation factor Tu; Ef-G: Elongation factor G; RP S12 & L31: Ribosomal proteins S12 and L31. Accession numbers of the datasets are available in the methods section.

The presence of a cytochrome *cbb*_3_ oxidase operon in Azoamicaceae symbionts suggests that like their free-living counterparts, symbiotic denitrifying bacteria also typically have the capacity to respire oxygen in addition to nitrogen oxides. However, oxygen respiration can be secondarily lost as demonstrated by the example of *Ca*. A. ciliaticola. In the case of Azoamicaceae symbionts, the presence and expression of a terminal oxidase raises the compelling possibility that Azoamicaceae respiratory endosymbionts respire oxygen and generate ATP for their host, instead of, or in addition to, host mitochondria.

### Evolutionary history of respiratory enzymes in *Azoamicaceae*

Given the importance of nitrate- and oxygen-respiring genes for the function of the Azoamicaceae respiratory endosymbionts, we used phylogenetic analyses to investigate the evolutionary history of those genes. We assume vertical inheritance when Azoamicaceae genes form a well-supported clade with their UBA6186 relatives, whereas genes that are absent from the entire UBA6186 group, are considered more likely to be horizontally acquired by the Azoamicaceae. All the core complexes of the electron transport chain (complex I, II, III and V) are monophyletic and appear to be vertically inherited from the UBA6186 clade (Figure 1, Extended Data Figure 3). In contrast, the cytochrome *cbb_3_*oxidase (complex IV) appears to be horizontally transferred from the Pseudomonadales, with the closest homologs present in Alcanivoraceae and Berkiellales (Figure 4a). The horizontal acquisition of *cbb_3_* oxidase by the Azoamicaceae is supported by the observation that none of the UBA6186 genomes, and only a single genome in the sister clade Berkiellales, contain genes for a cytochrome *cbb_3_* oxidase. Instead, both clades typically encode a cytochrome *caa_3_* oxidase and a cytochrome *bd*-type alternative oxidase, which are both absent from the five Azoamicaceae genomes (Figure 1). The key genes encoding for the core enzymes of the denitrification pathway (*narG*, *nirK*, *norB* and *nosZ*) also appear to be horizontally acquired, from four different bacterial donor lineages (Extended data figures 3-6). The four gene clusters are located at three distinct loci in the Azoamicaceae genomes, with the genes encoding for nitrite reductase and nitric oxide reductase adjacent to each other (Supplemental table S1). The two most closely related sequences to the Azoamicaceae *narG* gene, encoding for the nitrate reductase, are encoded in genomes of Gammaproteobacteria of the PIVX01 and JACQPQ01 orders. However, it appears that these Gammaproteobacteria likely obtained this gene horizontally from an alphaproteobacterial lineage (Extended Data Figure 3). The *nirK* gene, encoding for the nitrite reductase is most closely related to uncultured members of the *Ignavibacteraceae* family and was likely horizontally acquired from a member of this clade (Extended Data Figure 4). The provenance of the nitric oxide reductase and nitrous oxide reductase cannot be conclusively established from the phylogenetic analyses (Extended Data Figure 5 and 6), but the Azoamicaceae *norB* gene (encoding for nitric oxide reductase), and to a lesser extent the *nosZ* gene (encoding for nitrous oxide reductase) are related to those found in predatory bacteria of the Bdellovibrionota and Myxococcota phyla, allowing for the possibility that these taxa might have acted as intermediate carriers of these genes.

**Figure 4.**
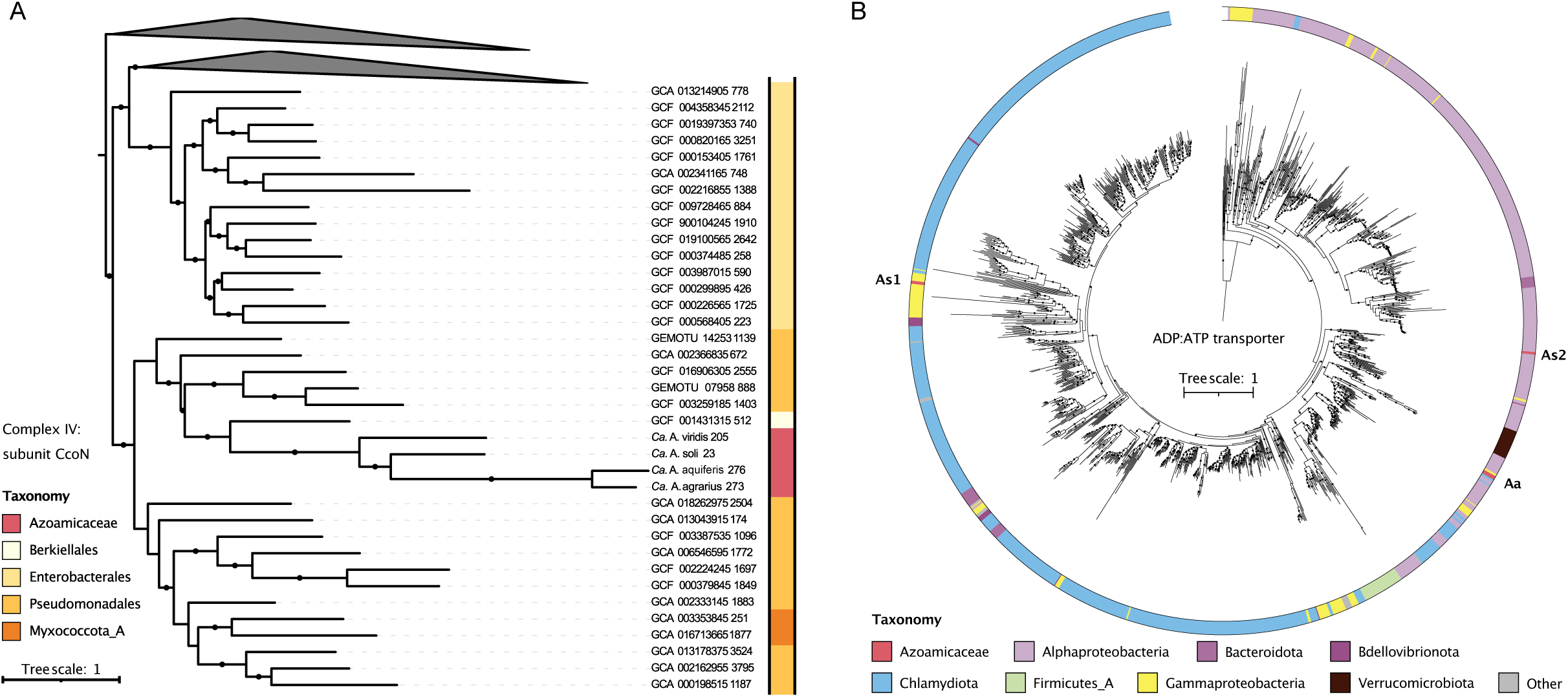
Phylogeny of key proteins in the Azoamicaceae. (A) Phylogenetic tree based on amino acid sequences of the catalytic subunit of the cbb_3_ cytochrome c oxidase (ccoN). Color strip indicates the GTDB assigned order within the class Gammaproteobacteria (or the phylum Myxococcota_A) of the respective genome containing the ccoN gene. (C) Phylogenetic tree based on amino acid sequences of the ATP/ADP transporter tlcA showing three independent origins for the Azoamicaceae genes. Both Azosocius genomes contain two copies (As1 & As2), and the three Azoamicus genomes contain third distinct tlcA copy (Aa). Color strip indicates the GTDB assigned taxonomy at the phylum or class (for Pseudomonadota) of the genome containing the tlcA gene. Sequences were obtained from genomes included in the genome taxonomy database and global catalog of earth’s microbiomes databases. Black circles on the branches indicate bootstrap values higher than 80% (A) or 90% (B).

Unlike the respiratory genes discussed above, the *tlcA* gene encoding an ATP/ADP transporter, is not monophyletic in the Azoamicaceae (Figure 4b) and seems to have been acquired by the Azoamicaceae or the larger UBA6186 order multiple times independently. Both Azosocius genomes contain one copy that was acquired by the UBA6186 order from a Chlamydiota organism and vertically inherited by Azosocius, but likely lost in Azoamicus. The Azosocius genomes contain a second copy likely directly acquired from a Alphaproteobacterium in the Rickettsiales order. The three Azoamicus genomes contain a third distinct tlcA copy, also likely originating from a Rickettsiales bacterium (Graf et al. 2021) and shared with one of the UBA6186 genomes, hinting at the possibility of vertical inheritance and loss in Azosocius (Figure 4b).

### Genetic potential for carbon metabolism and cofactor biosynthesis in Azoamicaceae

While the genes encoding the key enzymes of the respiratory pathway are highly conserved across the Azoamicaceae, there is less consistency regarding the distribution of genes involved in carbon metabolism across the four new genomes (Figure 2d). Three of the four new genomes encode genes for both the oxidation of malate to pyruvate (*maeA*) as well as to oxaloacetate (*mdh*). The *Ca*. A. ciliaticola genome only encodes *mdh*, which was highly transcribed *in situ*, leading to the suggestion that malate acts as the primary electron donor for denitrification (Graf et al. 2021). This is supported by the *Ca.* A. aquiferis transcription data analyzed here, showing that *mdh* is the third highest transcribed gene on average, across seven datasets. Conversely, no transcription of *maeA* could be detected in 6 out of 7 datasets (Figure 3, Supplemental table S5). Surprisingly, the smallest of the five genomes (*Ca.* A. soli) does not encode either of these enzymes for malate oxidation (Figure 2d), although it does retain the anion permease proposed to import malate in *Ca.* A. ciliaticola (*yflS*, Supplementary table S1) (Graf et al. 2021). From the present data, it can not be excluded that *Ca.* A. soli oxidizes malate using host-encoded malate dehydrogenase, but it is also possible that succinate is oxidized by complex II and used as an electron donor instead (Figure 2d).

Beyond the potential for malate and succinate oxidation, three of the genomes also encode 2-oxoacid:ferredoxin reductase (*korAB*) and succinyl-CoA synthetase (*sucCD*) that can oxidize 2-oxoglutarate to succinyl-CoA, and ultimately to succinate for further oxidation to fumarate using complex II. The *sucCD* genes are lacking in *Ca.* A. soli as well as in *Ca.* A aquiferis, the latter of which also appears to have lost the *korAB* genes (Figure 2d, Figure 5), suggesting use of only malate and (possibly) succinate as electron donors, or replacement of korAB and/or sucCD by host proteins.

**Figure 5.**
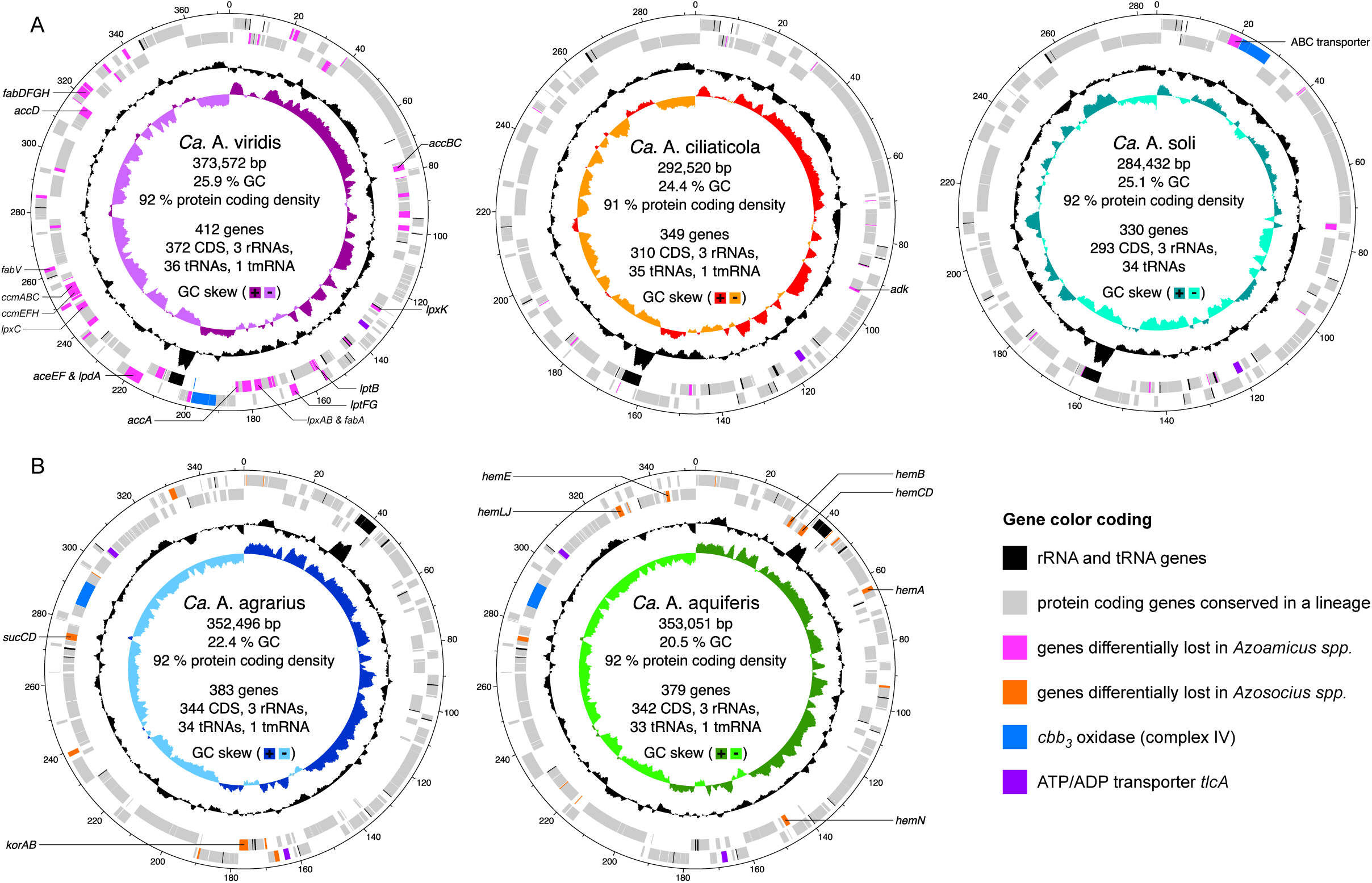
Differential gene loss in Azoamicaceae genomes. Circular genome maps of Ca. Azoamicus spp. (A) and Ca. Azosocius spp. (B) genomes, with rings (inside to outside) indicating GC skew, GC content, reverse strand genes, forward strand genes. rRNA and tRNA genes are indicated in black, protein coding genes encoded in multiple genomes of a genus are indicated in grey. Protein coding genes differentially lost in Azoamicus are indicated in pink, and differentially lost in Azosocius in orange. The cytochrome cbb_3_ oxidase and associated genes are highlighted in blue, and the ATP/ADP transporter genes in purple.

In addition to genes involved in respiration, other genes conserved in all 5 genomes include those involved in the biosynthesis of metal cofactors. These include the *sufBCDSU* genes for iron-sulfur cluster biosynthesis, as well as the *moaABCDE*, *moeAB*, and *mobA* genes for the assembly of the molybdopterin cofactor required for nitrate reductase (Figure 5). In contrast, the *modABC* genes encoding a transporter for molybdenum are lost from the two Azosocius genomes. Interestingly, a near complete gene set for heme biosynthesis (*hemABCDEJLN* and *glnS*) remains in only the *Ca.* A. aquiferis genome, although *hemH* is missing. On the other hand, the cytochrome maturation system encoded by *ccmABCEFH* was found only in *Ca.* A. viridis.

Notably, the genome of *Ca.* A. viridis also retained the capacity for fatty acid biosynthesis from pyruvate. Malonyl-CoA required for fatty acid biosynthesis can be formed from malate by the activities of malic enzyme (*maeA*), pyruvate dehydrogenase (*aceEF*, *lpdA*), and acetyl-CoA carboxylase (*accABCD*) (Figure 5). The latter two complexes are only retained in *Ca.* A. viridis. Fatty acids can then be synthesized from malonyl-CoA by the enzymes of the type II fatty acid biosynthesis pathway (*fabADFGHV*) (Figure 5) (Zhang and Rock 2008). However, the genes encoding the membrane proteins required to convert fatty acids to phosphatidic acid are missing from *Ca.* A. viridis. Apart from two genes (*cdsA*, *pgsA*) no other genes for synthesis of phosphatidylglycerol or phosphatidylethanolamine were found. Instead, it encodes the same Mla pathway thought to be used for lipid import in the other Azoamicaceae (Graf et al. 2021) (Supplementary table S1). *Ca.* A. viridis additionally encodes the inner membrane component of the LPS export system (*lptBGF*), as well as several of the steps needed to synthesize lipid-IVa (*lpxABCDK*). However, as the genome is lacking *lpxH,* a gene considered essential for biosynthesis of the Lipid A moiety, as well as key genes of the heptose biosynthesis pathway and the outer membrane component of the export system (*lptADE*), it seems unlikely that *Ca.* A. viridis can synthesize LPS for a typical outer cell membrane. However, it is peculiar that the LipidA biosynthesis pathway has been retained in the strongly reduced genome of Ca. A. viridis, suggesting that it may have a yet unidentified role in the symbiosis.

### Presence and distribution of Azoamicaceae in global amplicon datasets

In addition to comparative genomics, the newly recovered genomes also provide an opportunity to assess the distribution of Azoamicaceae respiratory endosymbionts in diverse ecosystems across the world. For this, we searched public 16S rRNA gene amplicon datasets for the presence of the five endosymbiont 16S rRNA gene sequences. We identified 998 unique datasets containing putative Azoamicaceae sequences (≥ 3 reads at ≥ 95 % sequence identity). These datasets encompass samples from all continents, suggesting a global distribution of Azoamicaceae endosymbionts (Figure 6). It is interesting to note, that despite all four of our new genomes being from groundwater, only 18 samples (2%) containing Azoamicaceae reads were annotated as ‘groundwater metagenomes’ in our global survey. However, the quality of sample type annotation of 16S rRNA gene amplicon datasets is highly variable, and samples of other categories (such as “freshwater metagenome” or “soil metagenome”) might encompass more groundwater samples. In any case, the two most common sample types in which putative Azoamicaceae sequences were found were aquatic/freshwater and wastewater/activated sludge (Supplemental table S4). The first described *Ca.* A. cilaticola genome was indeed recovered from anoxic lake water, and our analyses suggest that symbiont sequences most closely related (≥ 95 % identity) to *Ca.* A. ciliaticola are by far the most common and present in 808 of the 998 datasets (81 %). Relatives of *Ca.* A. agrarius were second most abundant and found in 143 datasets (14 %; Supplemental Table S4). The Azoamicaceae sequences were generally rare within the examined datasets, with Azoamicaceae reads accounting for ≦ 0.1 % of reads in 90 % of the retrieved datasets. This is consistent with the presence of endosymbionts constrained by host abundance, and may partly explain why the Azoamicaceae have only recently been discovered. Reads related to the Azoamicaceae accounted for ≥ 1 % of the reads in only 10 out of 998 datasets, which represent wastewater/activated sludge (6), freshwater lake (2), human gut (1), and groundwater (1). In general, our results indicate that wastewater is a widespread and probably important habitat for the symbionts that warrants future attention. Even though the five genomes currently known do not represent the full diversity of the Azoamicaceae, they already suggest a ubiquitous presence of the Azoamicaceae symbionts in diverse ecosystems around the world.

**Figure 6.**
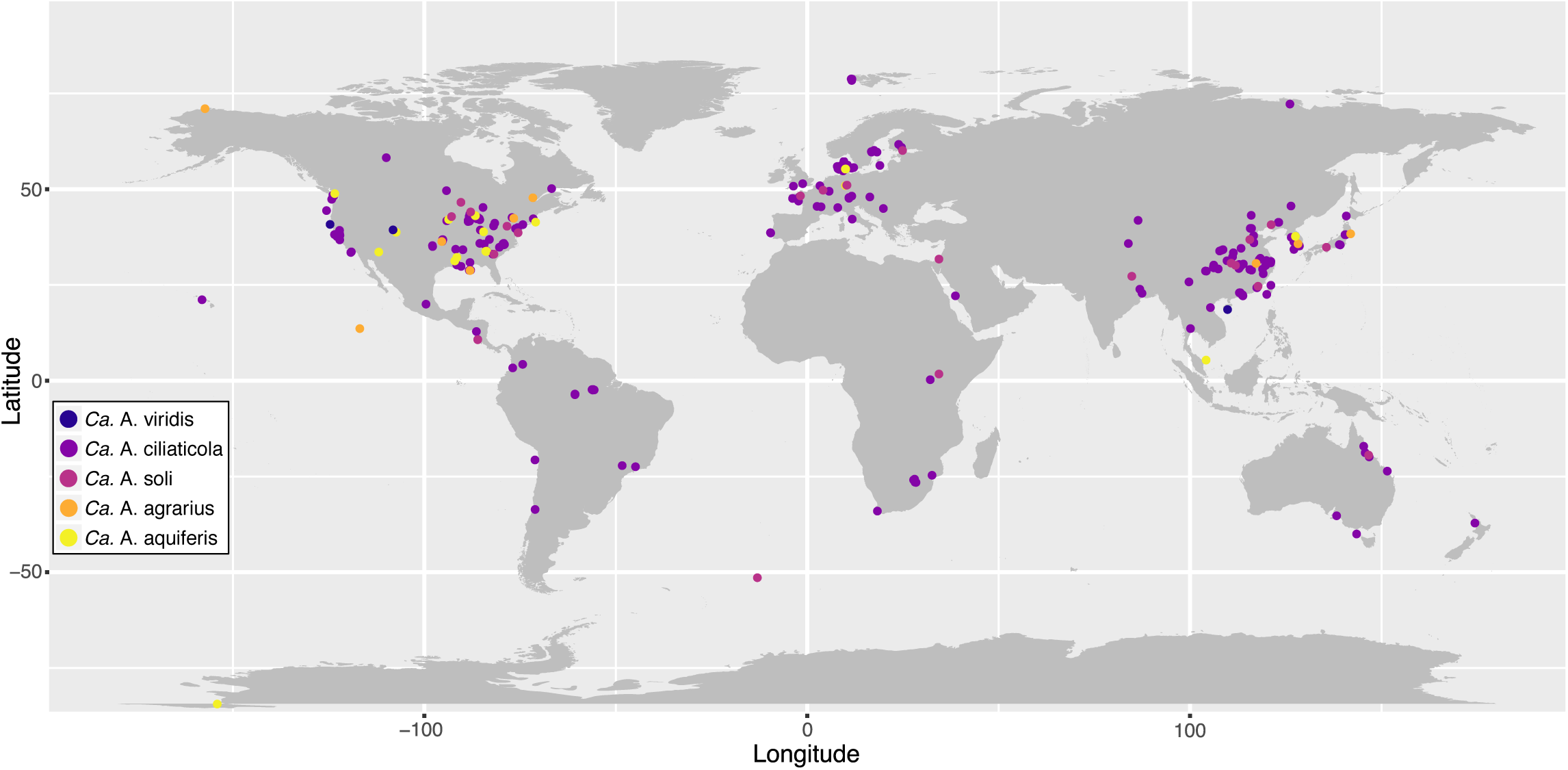
Detection of the Azoamicaceae in 16S rRNA gene amplicon sequencing datasets. Detection of the Azoamicaceae 16S rRNA gene sequences in publicly available 16S rRNA gene amplicon sequencing datasets shows a global distribution across all continents. Points represent amplicon datasets, and are colored according to their affiliation with the Azoamicaceae genome that have >95% 16S rRNA gene identity to reads in the dataset. The hotspots in the global north likely reflect sampling bias.

## Discussion

The four Azoamicaceae genomes described in this study greatly expand our understanding of the diversity and distribution of respiratory endosymbiosis. We show that Azoamicaceae are globally distributed, and have the genomic potential for both denitrification and aerobic respiration. Of the metabolic capacities encoded in the new endosymbiont genomes it is undoubtedly the capacity to respire oxygen that has the most profound ecological implications. The presence of cbb3 oxidase can explain the broad distribution of the symbiont in seasonally or permanently oxic environments, thus considerably expanding the potential niches for this symbiosis. Retention of the *cbb_3_* terminal oxidase in these extremely reduced genomes, as well as the comparatively high *ccoNOP* transcription in *Ca.* A. aquiferis, strongly implies an active role in endosymbiont and host physiology, by potentially conveying the capacity to respire oxygen to its host. While we cannot conclusively identify the host of the new symbiont lineages, 18S rRNA gene sequences that are related to the lacustrine host of *Ca.* A. ciliaticola could be found in the metagenomes containing *Ca.* A. agrarius and *Ca.* A. aquiferis. Additionally, *Ciliophora* were abundant in a molecular analysis of the wells of the Hainich CZE (Herrmann et al. 2020), further suggesting the *Ca.* A. aquiferis host is likely a ciliate.

Our comparative genomics of the five Azoamicaceae genomes shows a varying degree of genome erosion, as observed in insect HBEs (Moran, McLaughlin, and Sorek 2009; McCutcheon and Moran 2011). The Azoamicaceae seem to converge on a minimal gene set encoding the genes required for ATP generation using a denitrifying and oxygen respiring respiratory chain, with the conserved core genes comprising over 80% of the protein complement of the smallest genomes. The loss of biosynthetic pathways essential for the respiratory function, such as heme biosynthesis and cytochrome c maturation, indicates that host proteins are targeted to the endosymbiont. Host protein targeting has previously been observed in rare cases in insect HBEs (Nakabachi et al. 2014; Bublitz et al. 2019) and more extensively in the chromatophore of *Paulinella chromatophora* (Nowack and Grossman 2012). These observations have blurred the boundaries between HBEs and organelles, and the extant Azoamicaceae may represent another snapshot of the transition between the two. While comparative genomic analyses can provide a great insight into the evolutionary history and metabolic potential of these enigmatic endosymbionts, the establishment of an experimentally tractable laboratory culture of the host is essential to test the exciting hypotheses.

## Acknowledgments

We thank Falko Gutmann and Robert Lehmann for assistance with sampling the aquifers of the Hainich CZE as part of the Collaborative Research Centre AquaDiva (CRC 1076 AquaDiva - Project-ID 218627073) of the Friedrich Schiller University Jena, which is supported financially by the Deutsche Forschungsgemeinschaft.. This study used publicly available data from Bioprojects PRJNA640378, PRJEB36523, and PRJNA512237 and we thank the authors of those studies for making their data available. This study was financially supported by the Max Planck Gesellschaft.

## Online Methods

### Database searching and cMAG curation

The 16S rRNA sequence and nitrate reductase sequence of *Ca.* A. ciliaticola were used to search public genome databases. This led to the identification of the *Candidatus* Azosocius agrarius cMAG consisting of a single circular contig with 100 bp overlap (CP066692), binned from a groundwater sample taken in Modesto, CA, USA (37.661915 N, 121.114554 W) in 2018 (SAMN15459604) (He et al. 2021). For this study, we removed the overlap and started the contig at the putative origin of replication, resulting in a contig of 352,496 bp. The fasta file with the modified cMAG is provided as Supplemental File S1.

To identify further samples containing respiratory endosymbionts in groundwater, we used the 16S rRNA gene of *Ca.* A. agrarius to query publicly available 16S rRNA gene amplicon sequencing data sets using the web search tool IMNGS (Lagkouvardos et al. 2016). This search led to matches with >95% sequence identity in amplicon datasets from a groundwater aquifer located near Hainich national park in Germany, through the long term AquaDiva project (Supplemental Table S4). We screened the assembled metagenomes from the AquaDiva project (51.119338 N, 10.469198 W; Bioproject PRJEB36523) (Overholt et al. 2022) for contigs matching the *Ca.* A. agrarius cMAG using BLASTn (Altschul et al. 1990) (version 2.6.0) with a minimum length cutoff of 3000 bp. We iteratively mapped the trimmed reads from datasets ERR3858113, ERR3858114, ERR3858115 on the resulting 13 contigs using coverM (https://github.com/wwood/CoverM), followed by reassembly of the mapping reads using Spades (Bankevich et al. 2012) (version 3.15) with the –isolate flag, resulting in a circular contig of 353051 bp after three iterations, as confirmed by visualization in Bandage (Wick et al. 2015) (version 0.9.0). This circular contig represents the cMAG of *Candidatus* Azosocius aquiferis and is deposited to genbank under accession number NCBI BioSample SAMN39831648 and NCBI GenBank accession <Accession number pending>.

For further endosymbiont discovery we constructed a database of proteins encoded by *tlcA* gene, encoding the NTT transporter required for the exchange of ATP and ADP with the host, and screened sequencing runs using a blast score ratio (BSR) approach as previously described (Rasko, Myers, and Ravel 2005; Speth and Orphan 2018). We then queried the SRA for groundwater datasets, and selected large and medium scale bioprojects to screen. We screened a combined 256 sequencing runs from bioprojects PRJNA640378, PRJEB36505, PRJEB28738, PRJNA530103, PRJNA268031, PRJNA292723, PRJEB32173, PRJEB14718, PRJNA513876, and PRJNA512237 encompassing groundwater samples from Australia, Saudi Arabia, Germany, and USA (Colorado, Tennessee, California and Ohio). While we found evidence for the presence of putative endosymbionts (at low abundance) in several datasets, we only managed to reconstruct additional cMAGs, which was the quality threshold we set for this study, from sequencing runs from bioproject PRJNA512237 (Danczak et al. 2019). cMAG reconstruction for *Ca*. A. viridis and *Ca*. A. soli was performed by first co-assembling trimmed reads from datasets SRR8863434, SRR8863435, and SRR8863439 using MEGAHIT (D. Li et al. 2015) (version 1.2.9). The resulting contigs were searched using BLASTn (Altschul et al. 1990)(version 2.6.0) using the three other cMAGs as query, keeping hits with a minimum length cutoff of 3000 bp. This approach yielded 3 contigs, a circular contig representing the cMAG of *Candidatus* Azoamicus soli, and two contigs forming the MAG of *Candidatus* Azoamicus viridis. The *Ca.* A. viridis MAG was then circularized as described above for *Candidatus* A. aquiferis. The cMAG of *Candidatus* Azoamicus soli is available via NCBI BioSample SAMN39831885 and NCBI GenBank accession <Accession number pending> and the cMAG of *Candidatus* Azoamicus viridis is available via NCBI BioSample SAMN39831884 and NCBI GenBank accession <Accession number pending>.

### Genome annotation and comparative genomics

cMAGs were analyzed using anvi’o (Eren et al. 2021) (version 7.1), with gene calling by prodigal (Hyatt et al. 2010) (version 2.6.3). Annotation of protein coding genes was transferred from *Ca.* A. ciliaticola where appropriate based on bidirectional best hit DIAMOND (2.0.14.152) (Hernández-Salmerón and Moreno-Hagelsieb 2020), and the genes were annotated using KEGG (Kanehisa and Goto 2000), COG (Galperin et al. 2015) and PFAM (El-Gebali et al. 2019) annotation as integrated in anvi’o, followed by manual curation of the annotation. The final annotation is provided as Supplemental Table S1. The UpSet plot (Lex et al. 2014) of the cMAG gene content was generated from a presence absence matrix of the genes content using the UpSetR R package (Conway, Lex, and Gehlenborg 2017).

The average nucleotide identity between the cMAGs was calculated using fastANI (Jain et al. 2018) (version 1.33) with a fragment length of 1000 basepairs. Average amino acid identity was calculated using ezAAI (Kim, Park, and Chun 2021) (version 1.2.2) with default settings. 16S rRNA genes were extracted from the cMAGs using anvi’o (development version) (Eren et al. 2021), and pairwise identity was calculated using BLAST (Altschul et al. 1990).

Metabolic prediction of the cMAGs was done using the anvi-estimate-metabolism program (https://anvio.org/help/main/programs/anvi-estimate-metabolism/) as integrated in anvi’o (Eren et al. 2021) (development version) using KEGG modules annotation.

### Metaranscriptome analysis

The 18 metatranscriptome sequencing read datasets in BioProject PRJEB28738 were retrieved from NCBI and trimmed using the cutadapt (Martin 2011) wrapper trim-galore (https://github.com/FelixKrueger/TrimGalore) with default settings and automatic adapter detection. The trimmed reads were mapped against all gene sequences in the *Ca.* A. aquiferis genome using coverM (https://github.com/wwood/CoverM), with minimum identity 95 % and minimum length fraction 80 %, exporting the read counts per gene sequence. Based on total reads mapped, seven datasets were selected for analysis (Supplemental table S5). Read counts matching protein coding genes were used for the calculation of transcript per million (TPM) values (B. Li et al. 2010) and genes were ranked according to average TPM across the seven datasets. For visualization of the transcriptome in a heatmap (Figure 3), transcript abundance was recalculated as “transcript per thousand”, following the same procedure as for TPM but using a 10^3^ scaling factor instead of 10^6^ to account for the number of reads mapped to *Ca.* A. aquiferis. Datasets included in the heatmap are ERR2809165 (H52_1), ERR2809163 (H52_2), ERR2809153 (H52_3), ERR2809156 (H41_1), ERR2809155 (H41_2), ERR2809154 (H41_3), and ERR2809148 (H41_4). Samples H5-2_1, H5-2_2, and H5-2_3 represent replicated metatranscriptome data from the same well (H5-2) from sampling campaign PNK69. Sample H4-1_4 represents a metatranscriptomic data set from three pooled RNA preparations from well H4-1 from from sampling campaign PNK66.

### Global detection of Azoamicaceae

The 16S rRNA genes from the four novel cMAGs and the previously published *Candidatus* Azoamicus ciliaticola were used to screen 16S rRNA gene amplicon datasets for matches using the IMNGS web search tool (Lagkouvardos et al. 2016). 998 Datasets with more than 3 reads matching any of the five cMAGs at >= 95 % identity were retained for further analysis. The dataset IDs were used to retrieve sample metadata from NCBI, and sample coordinates were manually standardized. For 210 of the 998 datasets no coordinates were provided and these are therefore not included in Figure 6. To assess environmental distribution of the Azoamicaceae, the 998 datasets were grouped by their dataset description (defined upon dataset submission to INSDC databases).

### Putative host 18S reconstruction

To gain insight into potential host organisms for the organisms represented by the novel cMAGs, we attempted to reconstruct full length 18S rRNA gene sequences from their source datasets using Phyloflash (Gruber-Vodicka, Seah, and Pruesse 2020). No 18S rRNA gene sequences were retrieved from datasets SRR8863434, SRR8863435, and SRR8863439 (source data for *Ca.* A. viridis and *Ca.* A. soli). One 18S rRNA gene sequence was from each of ERR3858113, ERR3858114, ERR3858115 (source data for *Ca.* A. aquiferis). These three sequences were identical, and belonged to a ciliate of the Plagiopylea class. Four 18S rRNA gene sequences were retrieved from SRR12113421 (source data for *Ca.* A. agrarius). One of the four sequences obtained from SRR12113421 belonged to a ciliate of the Plagiopylea class, closely related to the sequences obtained from ERR3858113, ERR3858114, ERR3858115.

### Phylogenetic analyses

Phylogenetic analysis of protein coding genes (*nuoG, sdhA, qcrB, atpA, ccoN, narG, nirK, norB, nosZ, tlcA*) was done using protein sequences obtained from genomes in the genome taxonomy database (GTDB, version 207) (Parks et al. 2018) and the genomic catalog of Earth’s microbiomes (Nayfach et al. 2020). Sequences were aligned using MUSCLE (Edgar 2004) (version 3.8.31), and phylogenetic trees were calculated using IQ-tree (Nguyen et al. 2015) (version 2.2.0) with automatic model selection using ModelFinder (Kalyaanamoorthy et al. 2017) and ultrafast bootstrapping (1000 replicates) with UFBoot2 (Hoang et al. 2018).

Concatenated marker gene phylogenies were calculated based on protein sequences identified using hmmer (Eddy 1998) (version 3.3.2) with the Bacteria_71 HMM set included in Anvi’o (Eren et al. 2021) (version 7.1). The anvi-get-sequences-for-hmm-hits program was used to retrieve sequences for HMM hits, align them using MUSCLE (Edgar 2004) (version 3.8.31), concatenate alignments, and write a partition file. The concatenated alignments and corresponding partition files were used to calculate phylogenies using IQ-tree (Nguyen et al. 2015) with automatic model selection using ModelFinder (Kalyaanamoorthy et al. 2017) and ultrafast bootstrapping (1000 replicates) with UFBoot2 (Hoang et al. 2018).

## Etymology

### Description of ‘*Candidatus* Azoamicus viridis’ sp. nov

’*Candidatus* Azoamicus viridis’ (L. masc. adj. viridis, green pertaining to Greene county, where the sample containing the species was taken). A bacterial species identified by metagenomic analyses. This species includes all bacteria with genomes that show ≥95% average nucleotide identity to the type genome for the species to which is available via NCBI BioSample SAMN39831884 and NCBI GenBank accession <Accession number pending>.

### Description of ‘*Candidatus* Azoamicus soli’ sp. nov

’*Candidatus* Azoamicus soli’ (L. masc. n. soli, of the earth). A bacterial species identified by metagenomic analyses. This species includes all bacteria with genomes that show ≥95% average nucleotide identity to the type genome for the species to which is available via NCBI BioSample SAMN39831885 and NCBI GenBank accession <Accession number pending>.

### Description of *’Candidatus* Azosocius’ gen. nov

’*Candidatus* Azosocius’ (N.L. masc. n. Azosocius, combines the prefix azo-(N. L., pertaining to nitrogen) with socius (Latin, masc. n., associate); thus giving azosocius (‘associate that pertains to nitrogen’). A bacterial genus identified by metagenomic analyses and delineated according to Relative Evolutionary Distance by the Genome Taxonomy Database (GTDB). The type species of the genus is ‘*Candidatus* Azosocius agrarius’.

### Description of ‘*Candidatus* Azosocius agrarius’ sp. nov

’*Candidatus* Azosocius agrarius’ (L. masc. adj. agrarius, of the soil). A bacterial species identified by metagenomic analyses. This species includes all bacteria with genomes that show ≥95% average nucleotide identity to the type genome for the species to which is available via NCBI BioSample SAMN15435421 and NCBI GenBank accession GCA_016432505.1.

### Description of ‘*Candidatus* Azosocius aquiferis’ sp. nov

’*Candidatus* Azosocius aquiferis’ (N.L. gen. masc. n. aquiferis, of an aquifer). A bacterial species identified by metagenomic analyses. This species includes all bacteria with genomes that show ≥95% average nucleotide identity to the type genome for the species to which is available via NCBI BioSample SAMN39831648 and NCBI GenBank accession <Accession number pending>.

### Description of ‘*Candidatus* Azoamicaceae’ fam. nov

’*Candidatus* Azoamicaceae’ (A.zo.a.mi.ca.ce.ae. N.L. masc. n. Azoamicus. type genus of the family; N.L. suff. –aceae to denote a family; N.L. masc. pl. n. Azoamicaceae, the family of the genus Azoamicus). The description of the family ‘*Candidatus* Azoamicaceae’ is the same as that of the genus ‘*Candidatus* Azoamicus’. The type genus is ‘*Candidatus* Azoamicus’.

### Description of ‘*Candidatus* Azoamicales’ ord. nov

’*Candidatus* Azoamicales’ (A.zo.a.mi.ca.les. N.L. masc. n. Azoamicus. type genus of the order; N.L. suff. –ales to denote an order; N.L. masc. pl. n. Azoamicales, the order of the genus Azoamicus). A bacterial order identified by metagenomic analyses and delineated according to Relative Evolutionary Distance by the Genome Taxonomy Database (GTDB). Phylogenetic analyses in this work have shown that the previously published ‘*Candidatus* Azoamicus ciliaticola’ is assigned to an uncharacterized order with provisional designation UBA6186. Consequently, we propose to rename the UBA6186 order to ‘*Candidatus* Azoamicales’. the type genus of the order is ‘*Candidatus* Azoamicus’. The order is assigned to the class Gammaproteobacteria.

---

**Extended data table 1.** Genome features of the 5 Azoamicaceae genomes.

| **Organism name** | **Genome size** | **Genes** | **CDS** | **rRNA** | **tRNA** | **tmRNA** | **GC content (%)** | **coding density (%)** |
| --- | --- | --- | --- | --- | --- | --- | --- | --- |
| Azoamicus ciliaticola | 292520 | 349 | 310 | 3 | 35 | 1 | 24.4 | 91 |
| Azoamicus viridis | 373572 | 412 | 372 | 3 | 36 | 1 | 25.9 | 92 |
| Azoamicus soli | 284432 | 330 | 293 | 3 | 34 | 0 | 25.1 | 92 |
| Azosocius agrarius | 352496 | 383 | 344 | 3 | 34 | 1 | 22.4 | 92 |
| Azosocius aquiferis | 353051 | 379 | 342 | 3 | 33 | 1 | 20.5 | 92 |

**Extended data figure 1.**
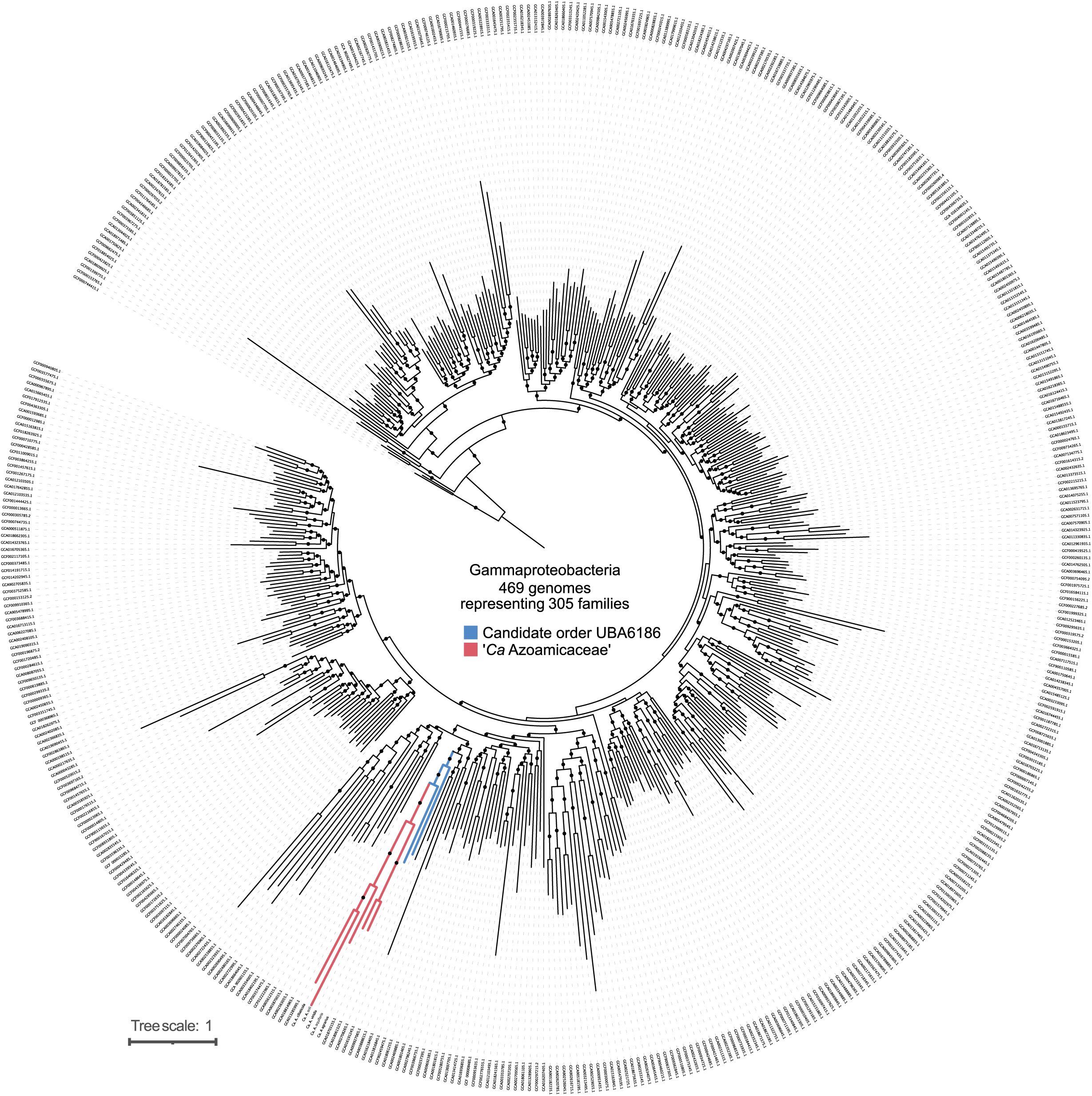
Phylogenetic placement of *Azoamicaceae* within the UBA6186 order in the Gammaproteobacteria class. Concatenated marker gene phylogeny of representatives of all 305 Gammaproteobacteria families in GTDB r207, including representatives of 2 genera per family if available. 146 families consisted of a single genus and are thus represented by a single genome. Genomes were selected based on type species status and contig number/completeness/redundancy.

**Extended data figure 2.**
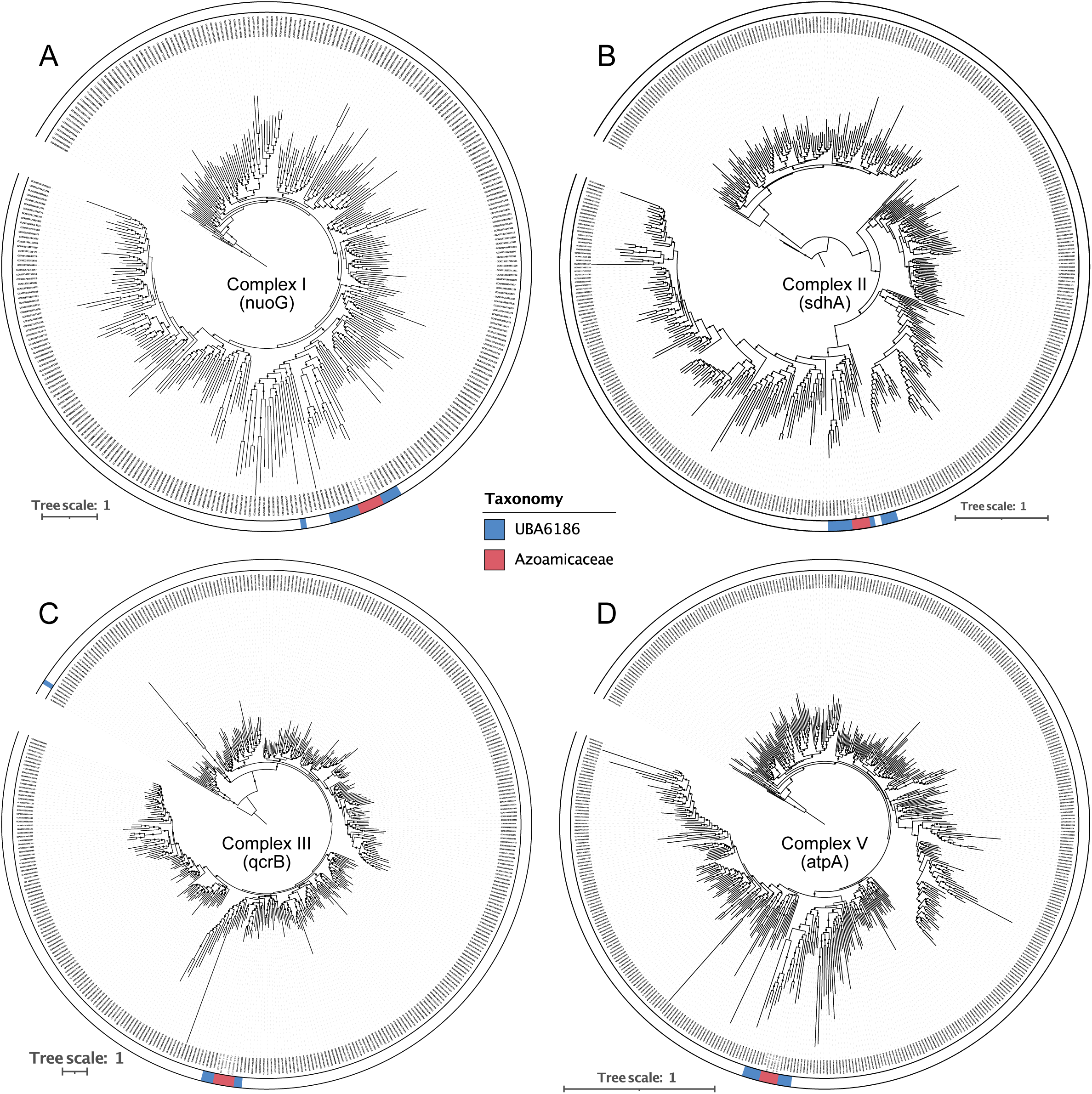
Respiratory chain complexes I, II, III and V are vertically inherited. Phylogenetic trees of the amino acid sequences of genes encoding subunits of A) complex I (*nuoG*), B) complex II (*sdhA*), C) complex III (*qcrB*), and D) complex V (*atpA*). Reference sequences were obtained from the 305 genomes included in Extended data figure 1, as well as all the UBA6186 genomes included in the genome taxonomy database (GTDB v207; GCA and GCF identifiers) and global catalog of earth’s microbiomes (GEM; GEMOTU identifiers) databases. Color strips indicate sequences from the Azoamicaceae family and UBA6186 order. Black circles on the branches indicate bootstrap values higher than 80%

**Extended data figure 3.**
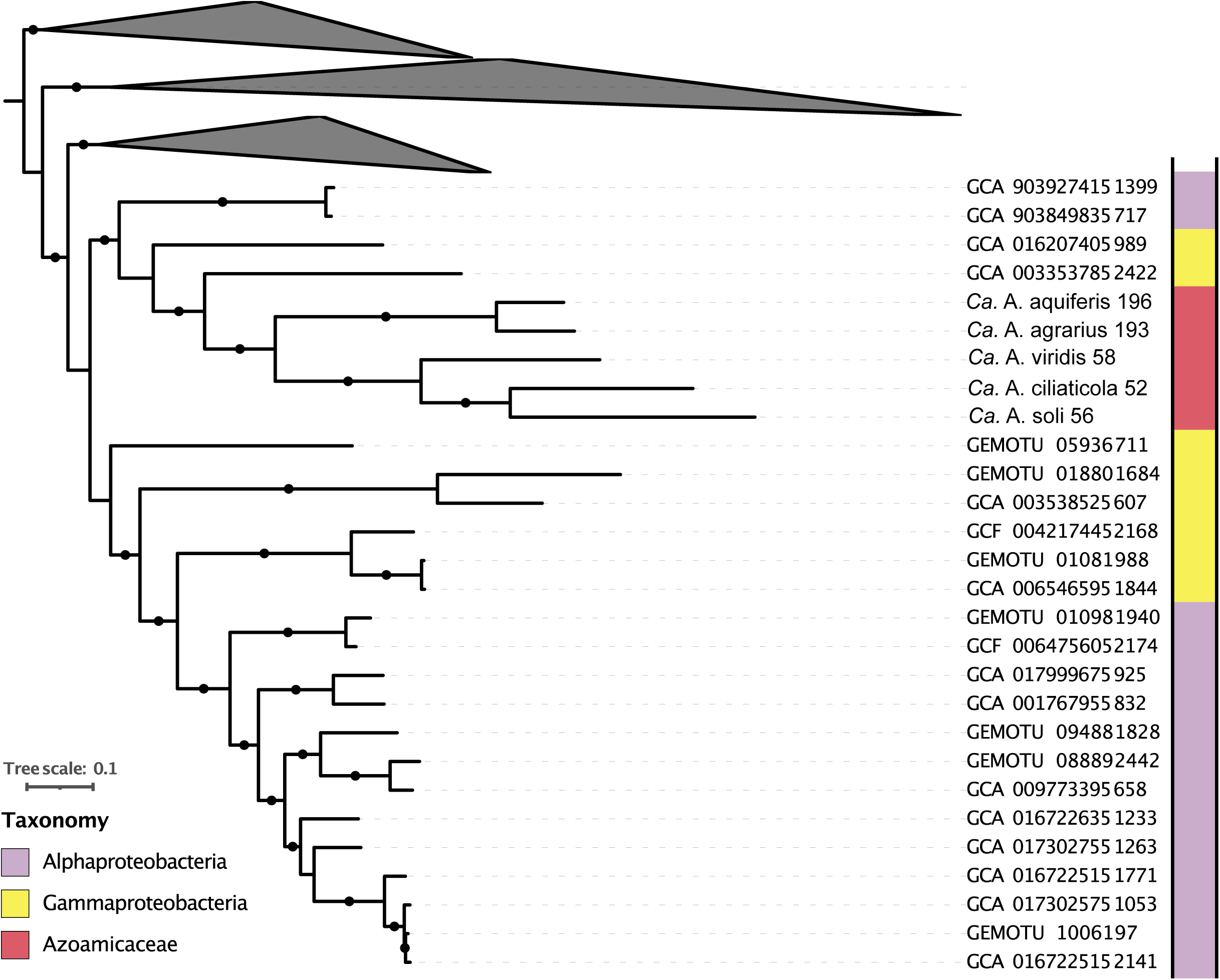
\**Nitrate reductase phylogeny suggests acquisition via lateral transfer.** Phylogenetic tree of the amino acid sequences of the molybdenum containing subunit of the nitrate reductase (narG). Sequences were obtained from genomes included in the genome taxonomy database (GTDB; GCA and GCF identifiers) and global catalog of earth’s microbiomes (GEM; GEMOTU identifiers) databases. Color strip indicates the GTDB assigned phylum (or class for Pseudomonadota) of the genome containing the *narG* gene, with colors consistent between Extended data figures 3-6. Black circles on the branches indicate bootstrap values higher than 80%.

**Extended data figure 4.**
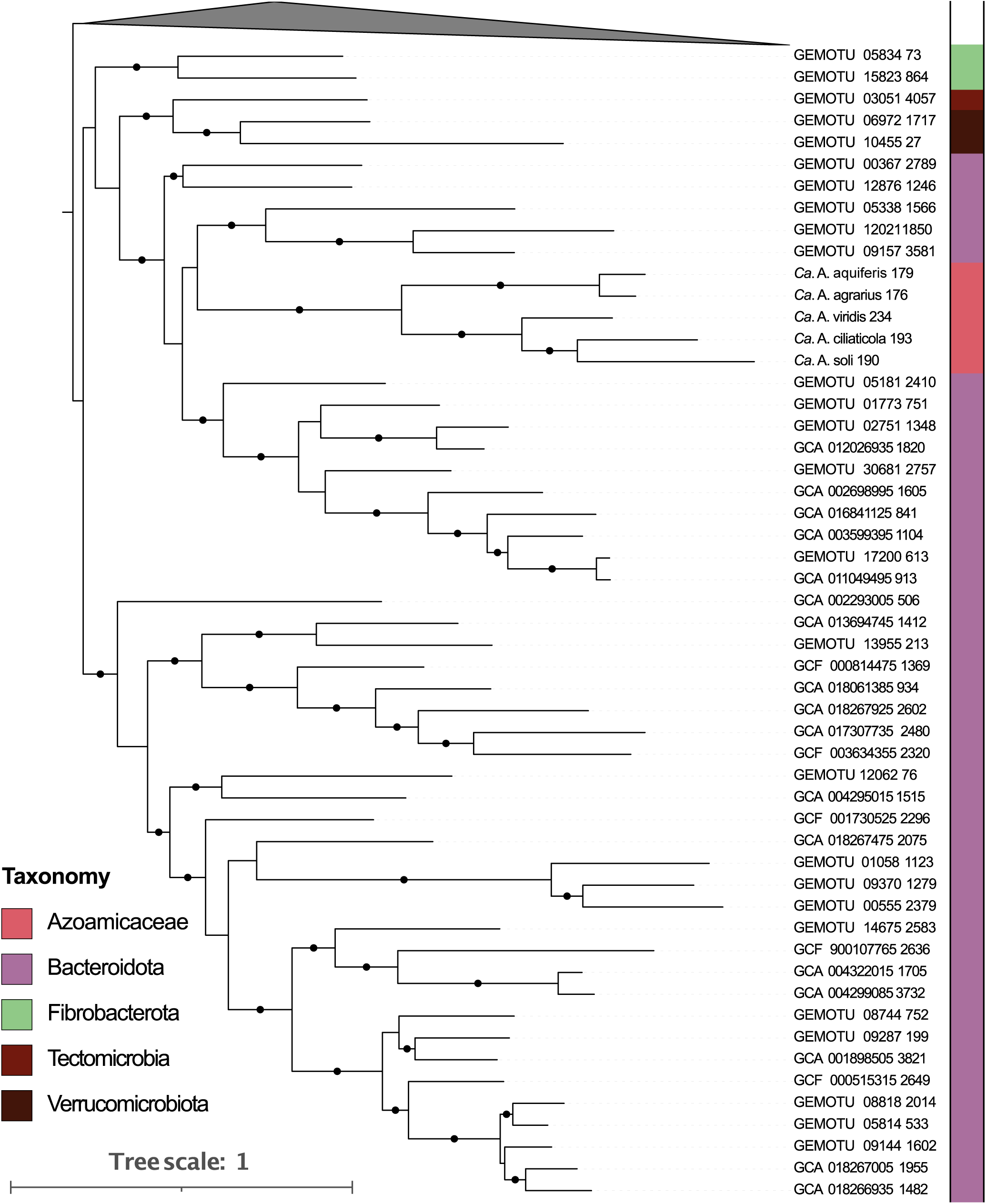
Nitrite reductase phylogeny suggests acquisition from Bacteroidota. Phylogenetic tree of the amino acid sequences of the copper containing nitrite reductase (nirK). Sequences were obtained from genomes included in the genome taxonomy database (GTDB; GCA and GCF identifiers) and global catalog of earth’s microbiomes (GEM; GEMOTU identifiers) databases. Color strip indicates the GTDB assigned phylum (or class for Proteobacteria) of the genome containing the *nirK* gene, with colors consistent between Extended data figures 3-6. Black circles on the branches indicate bootstrap values higher than 80%.

**Extended data figure 5.**
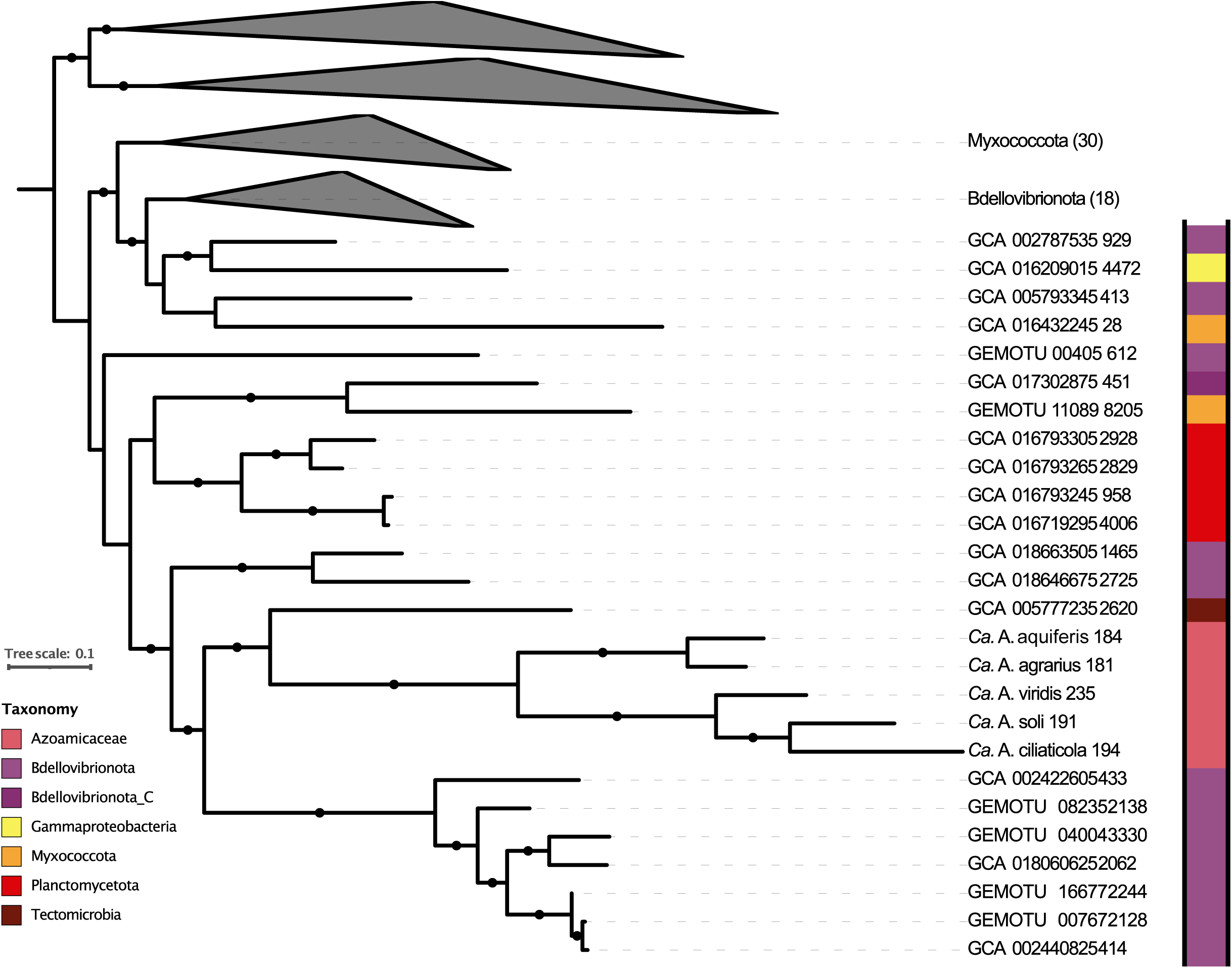
Nitric oxide reductase phylogeny. Phylogenetic tree of the amino acid sequences of the integral membrane subunit of the C-NOR type nitric oxide reductase (norB). Sequences were obtained from genomes included in the genome taxonomy database (GTDB; GCA and GCF identifiers) and global catalog of earth’s microbiomes (GEM; GEMOTU identifiers) databases. Color strip indicates the GTDB assigned phylum (or class for Proteobacteria) of the genome containing the *norB* gene, with colors consistent between Extended data figures 3-6. Black circles on the branches indicate bootstrap values higher than 70%.

**Extended data figure 6.**
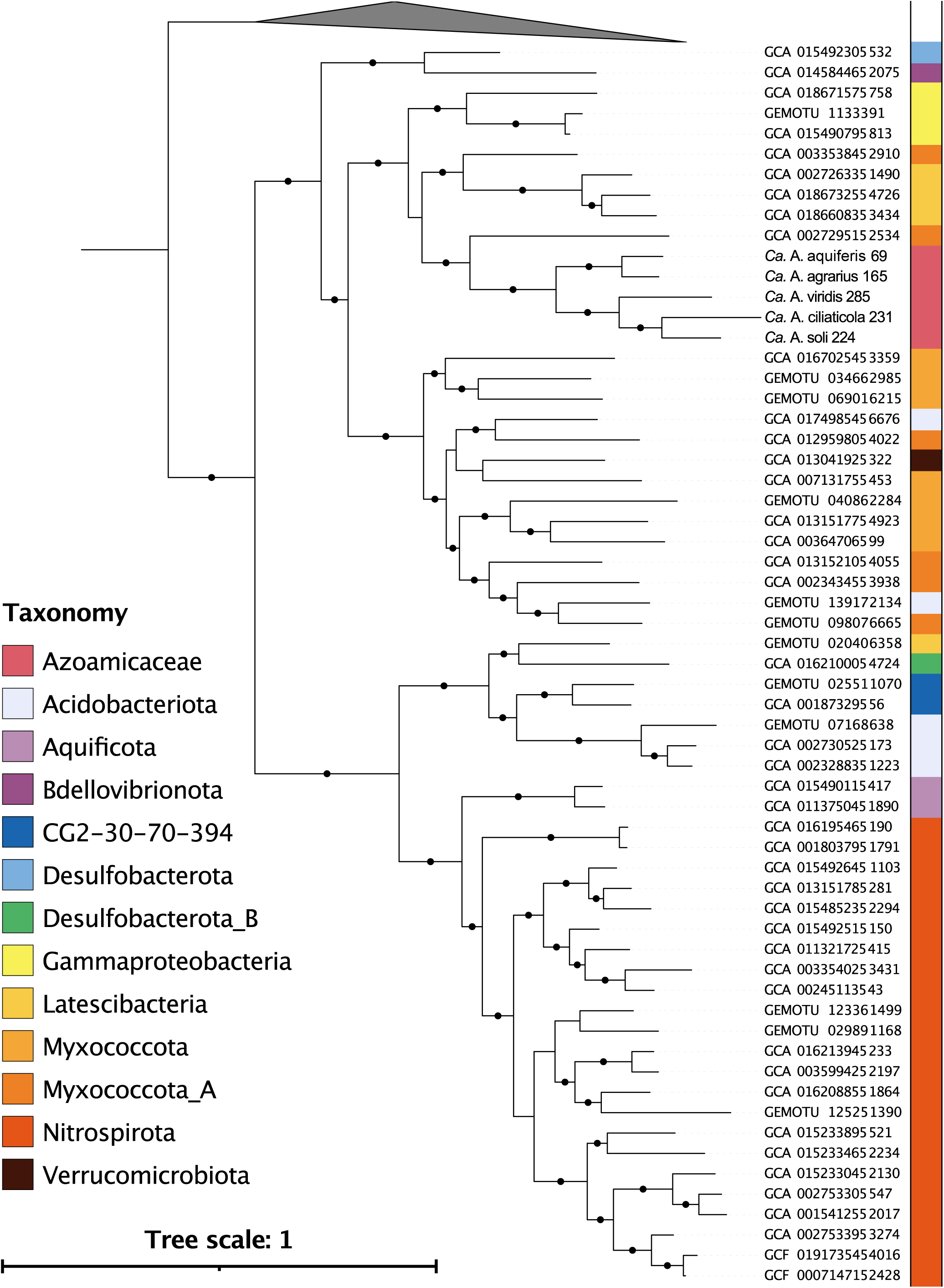
Phylogeny of nitrous oxide reductase suggests frequent transfers. Phylogenetic tree of the amino acid sequences of the catalytic subunit of nitrous oxide reductase (nosZ). Sequences were obtained from genomes included in the genome taxonomy database (GTDB; GCA and GCF identifiers) and global catalog of earth’s microbiomes (GEM; GEMOTU identifiers) databases. Color strip indicates the GTDB assigned phylum (or class for Proteobacteria) of the genome containing the *nosZ* gene, with colors consistent between Extended data figures 3-6. Black circles on the branches indicate bootstrap values higher than 80%.

