## Supplemental Text for "Genetic potential for aerobic respiration and denitrification in globally distributed respiratory endosymbionts"

**Affiliations:**
1 Department of Biogeochemistry, Max Planck Institute for Marine Microbiology, Bremen, Germany
2 Aquatic Geomicrobiology, Friedrich Schiller University, Jena, Germany

3 Cluster of Excellence Balance of the Microverse, Friedrich Schiller University, Jena, Germany

§ Present address: Division of Microbial Ecology, Centre for Microbiology and Environmental Systems Science, University of Vienna, Vienna, Austria

**Supplemental materials overview:**

Supplemental text - this document

Supplemental file S1 - reoriented Ca. Azosocius agrarius contig

Supplemental table S1 - Azoamicaceae cMAG gene annotation and presence/absence

Supplemental table S2 - Azoamicaceae cMAG metabolic predictions

Supplemental table S3 - Azoamicaceae cMAG and 16S rRNA gene similarity

Supplemental table S4 - Azoamicaceae IMNGS amplicon analysis results

Supplemental table S5 - *in situ* metatranscriptome analysis of Ca. Azosocius aquiferis

**Supplemental text**

UBA6186 order

The UBA6186 order is an uncultivated clade sister to the Berkiellales order, within the Gammaproteobacteria class. The clade was first defined in a large-scale assembly and binning effort of publicly available metagenomes that labeled the metagenome assembled genomes recovered as “Uncultivated Bacteria and Archaea (UBA)” (Parks et al. 2017). As of GTDB release 207, the UBA6186 clade consists of 17 MAGs, representing 9 species clusters (Parks et al. 2022). The existing UBA6186 MAGs were recovered from a diverse range of high latitude environments, namely oil sands tailings pond in Alberta, Canada (Tan et al. 2015), forest soils in Massachusetts, USA (Alteio et al. 2020), wastewater treatment plants in Germany (Schneider et al. 2021) and Canada (Spasov et al. 2020), the Saanich inlet at Vancouver island (Lin et al. 2021), several freshwater lakes with oxygen stratification in Europe and North America (Buck et al. 2021; Nayfach et al. 2020) and a saline lake in Antarctica (Nayfach et al. 2020). To the best of our knowledge, no experimental studies or comparative genomics analyses of the UBA6186 clade have been published, and our work defines *Ca.* Azoamicus as the type genus of the order.

Justification for the genus and species designation of the novel cMAGs

The five cMAGs clearly form two distinct lineages, with both cMAGs obtained from OHIO (designated *Candidatus* Azoamicus viridis and *Candidatus* Azoamicus soli) grouping with previously described *Candidatus* Azoamicus ciliaticola. The other two cMAGs, obtained from California and Germany and designated *Candidatus* Azosocius agrarius and *Candidatus* Azosocius aquiferis respectively, form a separate branch (Figure 1, Extended Data Figure 1). When comparing the cMAGs using common sequence similarity metrics (16S rRNA gene identity, average nucleotide identity (ANI), and average amino acid Identity (AAI), Supplemental table S4), it is clear that they represent distinct species. The maximum 16S identity, ANI, and AAI (between *Ca.* A. agrarius and *Ca.* A. aquiferis) are 94.2 %, 82.8 %, and 74.6 % respectively, all well below proposed species boundaries (Jain et al. 2018; Konstantinidis and Tiedje 2007) (Supplemental Table S3).

Furthermore, it is clear that *Ca.* A. agrarius and *Ca.* A. aquiferis belong to the same genus, as their 16S rRNA identity, ANI and AAI all fall in the range previously proposed for genera (Konstantinidis and Tiedje 2007; Barco et al. 2020). For the three cMAGs we designated Azoamicus, the genus designation is a little more complicated. Their pairwise ANI values are similar to the pairwise ANI values with either Azosocius cMAG, indicating that this metric has saturated. The pairwise AAI between the Azoamicus cMAGs (59.4 % - 60.8 %) is slightly higher than their pairwise identity with the Azosocius cMAGs (53.9 % - 55.2 %), as is their pairwise 16S rRNA identity (89.8 % - 92.3 % versus 86.7 % - 88.7 %), agreeing with the phylogenetic analyses (Supplemental Table S3, Figure 1, Extended Data Figure 1). However, the latter two metrics even fall below proposed cutoffs for genus level.

We choose to propose that each of these two lineages represent a single genus, as it is likely that the cMAGs represent fast evolving genomes and thus that the sequence similarity metrics likely overestimate their evolutionary distance. We also noticed that there is extensive synteny between the cMAGs within a lineage, but not between the cMAGs from two different lineages (Supplemental Table S1). We thus propose that conservation of genome structure should be used in addition to sequence similarity metrics to delineate the genus boundaries of these putative endosymbionts.

Description of gene content

As discussed in the main text, 297 genes in the Azoamicaceae pangenome are conserved between the two lineages. In addition to the genes shared between the two lineages, there are 172 genes unique to either Azoamicus or Azosocius (Figure 2b, supplemental table S1). Of these, 107 genes are present only in Azoamicus genome(s), with the majority (57) unique to the largest *Ca.* A. viridis genome, which contains a full gene complement to synthesize lipids from pyruvate (Supplemental table S1). In addition, *Ca.* A. viridis exclusively encodes a system I type cytochrome c maturation pathway. Other Azoamicus-specific genes of note are protein trafficking genes encoding the signal recognition particle protein (*ffh*), its receptor (*ftsY*), and several genes in the outer membrane assembly complex (*bamBDE*). However, the gene encoding central subunit *bamA* was present in both lineages, lending support for the retention of an outer membrane in these endosymbionts, as previously proposed (Graf et al. 2021). Surprisingly, the genes encoding the molybdenum transporter (*modABC*), required for nitrate reductase, are also unique to the Azoamicus genomes. The 66 genes unique to Azosocius include the pathway for heme biosynthesis from glutamyl-tRNA that is present only in *Ca.* A. aquiferis, although the gene for the final reduction reaction (*hemH*) is missing. In addition, notable genes present only on both Azosocius genomes are a second copy of the ADP:ATP translocase required for ATP delivery to the host, as well as the gene for DNA mismatch repair protein mutH.
